# Effect of caldesmon mutations in the development of zebrafish embryos

**DOI:** 10.1101/2023.03.28.534118

**Authors:** Verneri Virtanen, Kreetta Paunu, Saana Niva, Maria Sundvall, Ilkka Paatero

**Affiliations:** Cancer Research Unit, Institute of Biomedicine, and FICAN West Cancer Center Laboratory, University of Turku, and Turku University Hospital, Kiinamyllynkatu 10, 20520 Turku, Finland; Department of Oncology, Turku University Hospital, PL52, 20521 Turku, Finland; Turku Bioscience Centre, University of Turku and Åbo Akademi University, Tykistökatu 6, 20520, Turku, Finland

**Keywords:** caldesmon, zebrafish, mutation, developmental biology

## Abstract

Cancer is a profound medical concern and better treatments are needed for cancer patients. Therefore, new cancer targets are constantly being studied. These targets need not only be relevant for cancer progression, but their modulation needs to be tolerated reasonably well by the host. Caldesmon is one of these proposed novel targets for cancer therapy. Therefore, we analysed effects of caldesmon mutations in normal development using genetically modified zebrafish embryos. We analysed mutations in both zebrafish caldesmon genes, *cald1a* and *cald1b* and analysed effects of either mutation alone or as in combination in double homozygous embryos using molecular, morphological and functional analyses. The effects of caldesmon mutations were mild and the gross development of zebrafish embryos was normal. The caldesmon mutant embryos had, however, alterations in response to light-stimulus in behavioral assays. Taken together, the effects of caldesmon mutations in the development of zebrafish embryos were reasonably well tolerated and did not indicate significant concerns for caldesmon being a potential target for cancer therapy.

## Introduction

Cancer is a one of the leading causes of death and the need for new therapeutics and therapeutic targets is evident. Many factors related to actomyosin contractions have been implicated as integral regulators of cell migration. In the directionally migrating non-muscle cells, actomyosin contractions produce motility via stress fibers, which can both rupture and strengthen focal adhesions by contraction [1]. Thus, factors interacting with the actomyosin bundles contribute to generating forces that migrating cells exert towards the extracellular matrix. The ability to migrate can be attained by cancer cells that undergo epithelial to mesenchymal transition (EMT), which entails the loss of polarity and cell-to-cell adhesions [2]. The multistage metastatic process set in motion by cancer cells invading into the adjacent tissues is thus initiated by factors promoting cell motility and EMT [3].

One of the factors in actomyosin pathway is caldesmon, which is encoded by the *CALD1* gene in humans [4]. The human *CALD1* produces two major isoforms by alternative splicing, h-caldesmon and l-caldesmon, which both share common actin-, myosin-, tropomyosin- and calmodulin-binding domains, but are exclusively expressed in muscle cells and in non-muscle cells, respectively [5,6]. Observations from of the role of *CALD1* in tumor invasion and metastasis [7–10], indicated the potential of *CALD1* as a therapeutic target for anti-cancer treatments.

In an optimal case, a potential therapeutic target has significant adverse effects only in tumor but not in healthy tissues. Finding and developing novel chemical compounds for new target molecules is slow and costly process [11] and many novel therapeutics fail due to safety concerns [12]. These failures may be related to toxic side effects or target-related toxicities. Already in the early stages of target validation, the potential of target-related toxicities can be evaluated by using gene-modified animal models[13]. Zebrafish are an affordable model, large number of mutant alleles are available, and the development of embryo and larvae can be easily analyzed [14]. Here, we chose to utilize mutant zebrafish models to characterize potential side effects of targeting *cald1*. We generated single-mutant zebrafish for both *cald1a* and *cald1b* genes as well as double-mutant zebrafish, which we then assessed for changes in morphology and in behavior during early development.

## Materials and Methods

### Zebrafish husbandry

Analyses of zebrafish embryos were carried out under the licenses MMM/465/712–93 (issued by the Finnish Ministry of Agriculture and Forestry) and ESAVI/9339/04.10.07/2016 and ESAVI/31414/2020 (granted by Project Authorization Board of Regional State Administrative Agency for Southern Finland) according to the regulations of the Finnish Act on Animal Experimentation (62/2006). The study was carried out in compliance with the ARRIVE guidelines.

### Zebrafish cald1a and cald1b mutants and genotyping

Zebrafish lines carrying *cald1a*^sa16974^ (*cald1a*^*-/-*^) and *cald1b*^sa24667^ (*cald1b*^*-/-*^) mutant alleles were obtained from European Zebrafish Resource Centre, Karlsruhe, Germany, and provided by Dr. Derek Stemple, Wellcome Trust Sanger Institute, Genome Research Limited, Hinxton, UK. The breeding stocks of these fish were kept as heterozygotes and intercrossed to generate homozygous embryos for the analyses. Zebrafish embryo DNA was extracted either using NucleoSpin TriPrep RNA, DNA, and protein extraction kit (Macherey-Nagel, Allentown, PA) (gene expression studies) or by using alkaline lysis (morphological and behavioural studies) dissolving the embryos in 50 mM NaOH 95 °C for 10 min. After alkaline lysis, 1 M Tris HCl pH 8 was used to neutralize the DNA solution. Genotyping PCR were done by using KASP genotyping assay (Biosearch Technologies, Hoddesdon, United Kingdom). A similar workflow was used in genotyping adult carrier fish.

### Morphological analysis

The embryos were anesthetized and imaged using Nikon Eclipse Ti2 (Nikon). Data from images was extracted in ImageJ. Overall morphology was evaluated visually. Eye size, head size, body length and pericardium size were measured with ImageJ. All measurements were done independently by two investigators. Although formal blinding and randomization was not carried out, the measurements and analyses were essentially blinded as samples were genotyped after experimentation and phenotyping.

### Quantitative Real-Time PCR

The RNA was extracted from zebrafish embryos using NucleoSpin TriPrep RNA, DNA, and protein extraction kit (Macherey-Nagel). RNA was further purified by using RNA Clean & Concentrator-25 kit (Zymo Research, Irvine, CA). After purification, RNA was reverse-transcribed with High-Capacity cDNA Reverse Transcription Kit (Thermo Fisher Scientific), and the cDNA was amplified with gene-specific primer pairs (*cald1a:* 5’ CACTTCGTTTGCCTCATCGC 3’, 5’ CGCCGATATGCCATCCTCTC 3’ and *cald1b*: 5’ CAGGAGGAAACAGTGCCAGA 3’, 5’ TCTTGCGGCTTTGTTGACAC 3’) using SYBR™ Green PCR Master Mix (Applied Biosystems, Bedford, MA). The quantities measured by real-time PCR were normalized to the *Rpl13* (5’GGCGGACCGATTCAATAAGGTTCTGATCATTG 3’, 3’CCAGAGATGTTGATACCCTCACACCTCAC 5’) expression level in each sample.

### Behaviour assays

Zebrafish behavioural assays were carried out using DanioVision (Noldus IT) instrument. Four 4dpf zebrafish embryos were placed in 96-well square bottom Whatman Uniplate multiwell plates (Sigma-Aldrich) in 300µl of E3 medium. The plate was transferred to prewarmed DanioVision instrument (28.5°C). After 30min adaptation phase in darkness, the embryos were subjected to three cycles of 5min light followed by 5min of darkness. After experimentation, the embryos were anesthesized with 200mg/l tricaine and DNA was extracted for genotyping. The data analysis was carried out using EthoVisio XT and GraphPad Prism 9. Prior to statistical analysis, the first 20mins were removed (adaptation phase), then baseline was calculated from 20-30min time points, and it was subtracted from values to correct for potential differences in baseline values. 2-way ANOVA with Holm-Sidak post-hoc test for multiple testing correction was carried out. To increase statistical power, the different genotypes were pooled. *wt* (n=52) = *wt* + *cald1a*^+/-^ + *cald1b*^+/-^ ; cald1a (n=24) = *cald1a*^-/-^ (*cald1b* either +/+ or +/-) ; *cald1b* (n=38) = *cadl1b*^-/-^ (*cald1a* either +/+ or +/-) ; *cald1a, cald1b* (n=13) = homozygous mutant for both genes. In some experiments, the embryos were stimulated with 20mM pentylene tetrazolium (PTZ), which was followed by similar adaptation and light-dark cycles. In these experiments neither pooling nor baseline subtraction was carried out.

## Results

### The in silico analysis of cald1a and cald1b mRNA expression

Zebrafish possess two caldesmon genes *cald1a* (ZFIN code: ZDB-GENE-030131-1629) located in chromosome 4 and *cald1b* (ZDB-GENE-090313-229) located in chromosome 25. To get insights into the potential role of caldesmon (*cald1a* and *cald1b*) in the developing zebrafish, we carried out in silico analysis of gene expression using published RNA-seq data set [15] using Gene Expression Atlas [16]. Both *cald1a* and *cald1b* were expressed during the development of zebrafish embryos (Fig 1A, whole embryo RNA-seq data). The *cald1a* was expressed throughout development, with lower levels during earlier time points and with weak maternal contribution. Expression of *cald1b* was low in the earlier stages and showed weak activation during gastrulation, whereas more robust expression was evident only from protruding mouth stage (72 hpf) onwards. Besides being expressed during the development, the temporal whole animal expression patterns did not yield profound insights into role of *cald1* during development.

**Figure 1.**
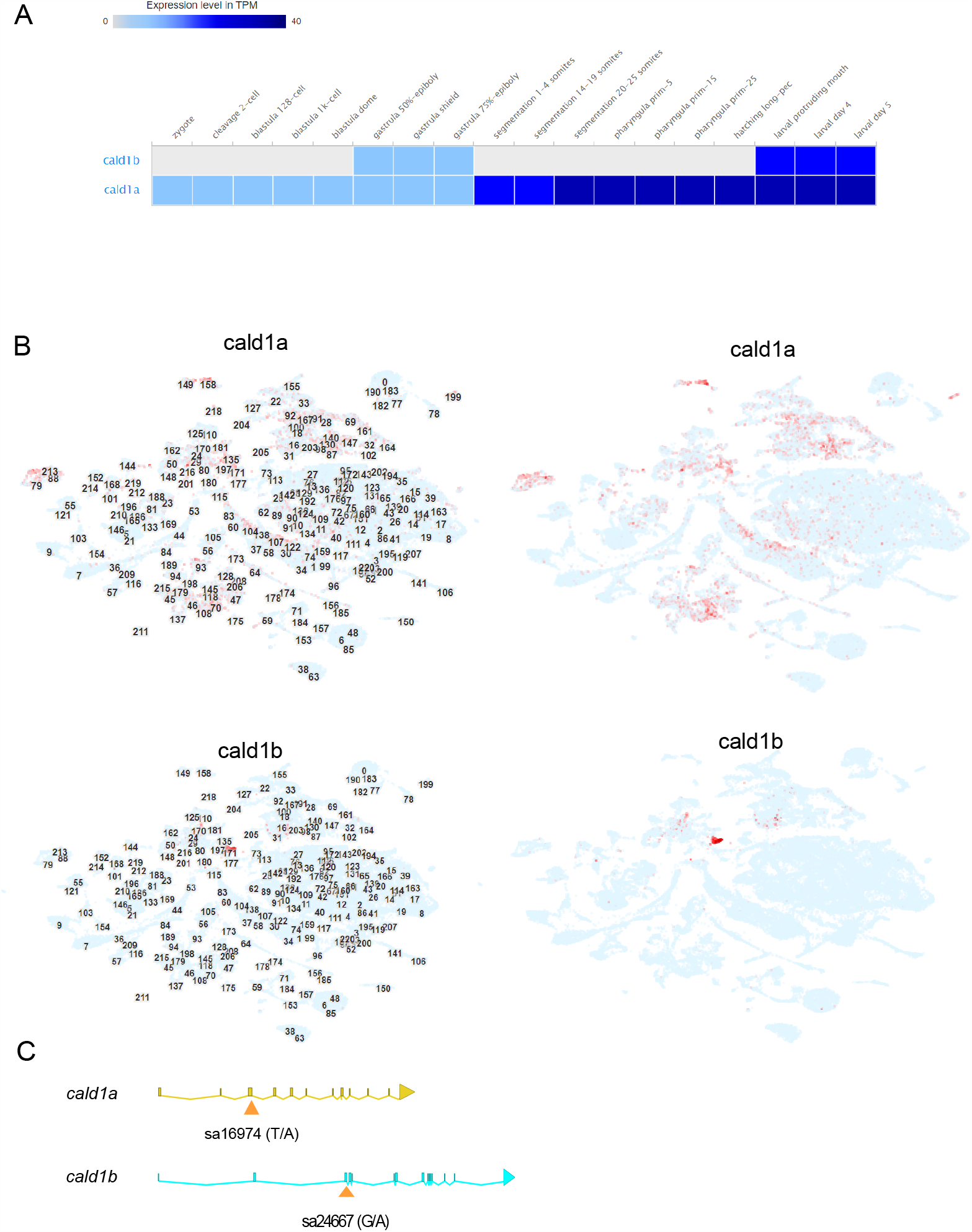
The *in silico* analysis of cald1a and cald1b mRNA expression, and the mutations. (A) Whole-embryo RNA-seq data for *cald1a* and *cald1b* from zebrafish embryo developmental time series. Data obtained from Expression Atlas (www.ebi.ac.uk/gxa/home, dataset E-ERAD-475, data downloaded January 3^rd^, 2023). (B) Single-cell RNA-seq data for developing zebrafish embryos. Data obtained from USCS Cell Browser (http://cells.ucsc.edu/?ds=zebrafish-dev, data obtained January 3^rd^, 2023). The numbers in left panels refer to cell cluster ID numbers and in the right panels these are omitted for clarity. (C) Schematic illustration of the location of mutations in *cald1a* and *cald1b* genes. Exons are marked as bars and connected by lines representing introns. Mutation sa24667 locates in exon 3 of *cald1a*, disrupting the essential splice site. The first 67 amino acids of 778 amino acids in *cald1a* are predicted to be intact in *cald1a*(sa24667) mutant. Mutation sa16974 locates in exon 3 and creates a premature stop codon. The first 190 amino acids of 778 amino acids in *cald1b* are predicted to be intact in *cald1b*(sa16974) mutant.

Recent advances in single-cell RNA-seq technology has enabled transcriptomic analysis of whole zebrafish embryos and single-cell level. We utilized pre-existing data set [17] using USCS Cell Browser [18] to analyze the spatio-temporal expression of cald1 transcripts during the development. *cald1a* was expressed broadly and especially in clusters associated with vasculature (clusters 79,88, 171, 213), heart (clusters 130, 147), neural crest (eg. 64, 87, 181), retina (30, 34, 58, 60, 83, 104, 138, 107, 122) and basal cells (47,145,70,108,118), whereas expression of *cald1b* was predominant in vascular smooth muscle cells (cluster 171) and neural crest (24,29, 31,170) (Fig.1B).

To study caldesmon in vasculature development we searched for potentially existing *cald1a* and *cald1b* mutant fish lines. Indeed, suitable mutant alleles, *cald1a*^sa16974^ (named here as *cald1a*^-/-^) and *cald1b*^sa24667^ (named here as *cald1b*^-/-^), were generated in Zebrafish Mutation Project [19] (Fig. 1C) and they were available through European Zebrafish Resource Centre EZRC. Therefore, we used these mutant lines for analysis of the role of *cald1a* and *cald1b* in developing zebrafish.

### No clear morphological phenotype for cald1a or cald1b single-mutant embryos

To investigate the effect of *cald1a* and *cald1b* on the morphology of developing zebrafish embryos (Fig 2A and B), we measured standard length, area of the cephalic region, eye area, and pericardial area from brightfield images of *cald1a* or *cald1b* mutant embryos (Fig 2C). To our surprise, no significant difference between the genotypes was detected in the body length, head size, eye size, or pericardium (Fig 2D–K).

**Figure 2.**
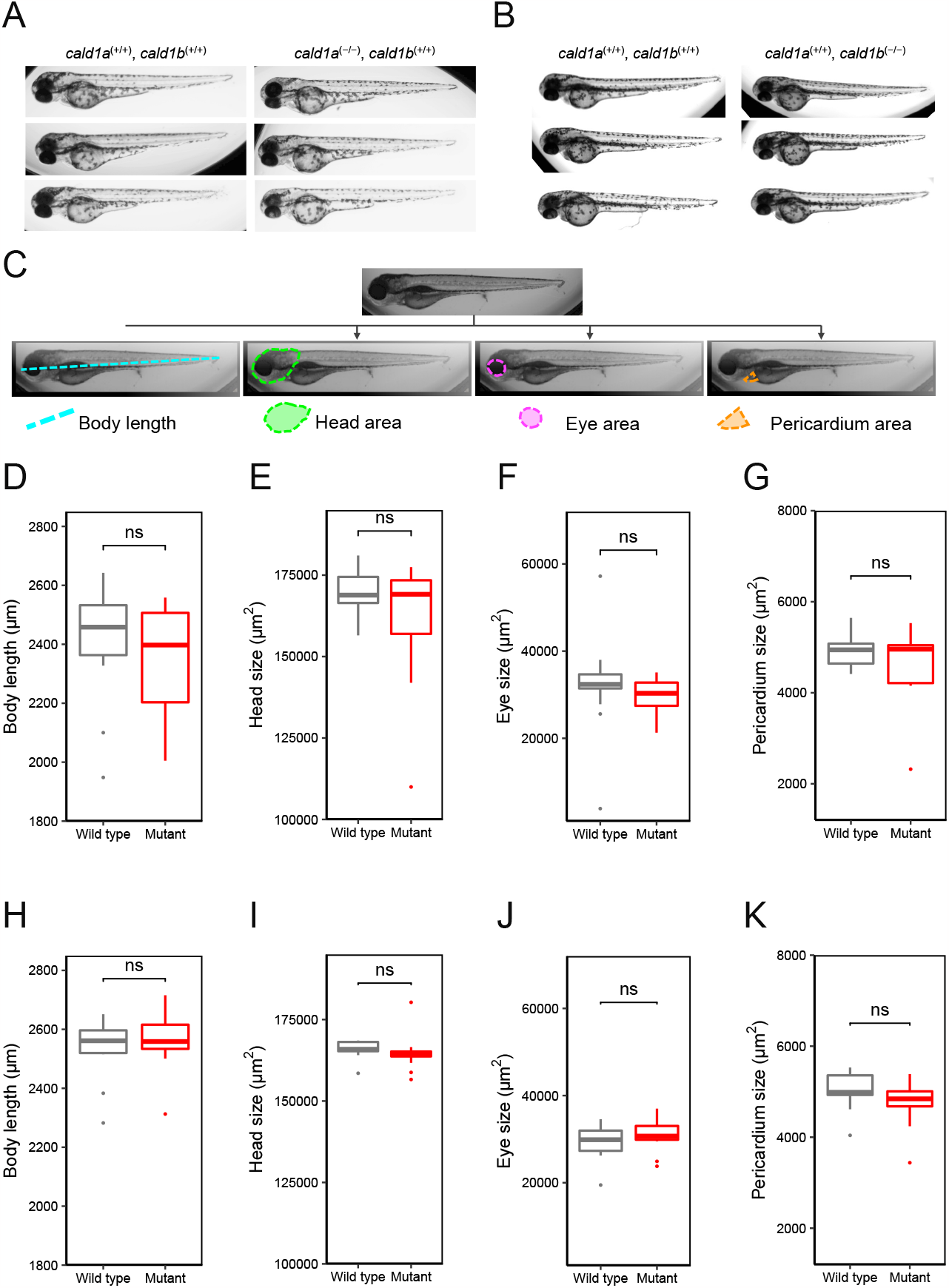
Analyses of the anatomy of *cald1a* and *cald1b* single-mutant zebrafish. (A) Representative brightfield images of wild-type zebrafish larvae *cald1a* (+/+), *cald1b* (+/+) (left) and single-mutant zebrafish larvae *cald1a* (−/−), *cald1b* (+/+) (right) 4 dpf. (B) Representative brightfield images of wild-type zebrafish larvae *cald1a (+/+), cald1b (+/+)* (left) and single-mutant zebrafish larvae *cald1a* (+/+), *cald1b* (−/−) (right) 4 dpf. 19) (C) Schematic illustration of the measurements taken from the zebrafish larvae. Body length, head area, eye area and pericardium area are highlighted over the duplicates of the original image with teal, green, purple, and orange, respectively. (D–G) Morphological measurements and analyses display no statistically significant changes in body length, head size, eye size, and in pericardial area when compared wild-type *cald1a* (+/+), *cald1b* (+/+) (n=22) larvae to single-mutant larvae *cald1a* (−/−), *cald1b* (+/+) (n=12). All measurements were done by two independent investigators. (H–K) Morphological measurements and analyses display no statistically significant changes in body length, head size, eye size, and in pericardial area when compared wild-type *cald1a* (+/+), *cald1b* (+/+) larvae (n=13) to single-mutant larvae *cald1a* (+/+), *cald1b* (−/−)(n=19). All measurements were done by two independent investigators.

### Regulation of cald1a and cald1b mRNA expression

To analyze *cald1a* and *cald1b* gene expression in mutant embryos we carried out qPCR analyses. In these qPCR analyses, *cald1a* mRNA was significantly reduced in the *cald1a*-mutant zebrafish larvae compared with the wild type (Fig. 3A). This was indicative of nonsense-mediated decay of mutated dysfunctional mRNAs [20]. Interestingly, *cald1b* mRNA expression was strongly increased in *cald1b*-mutant zebrafish larvae (Fig. 3B). This is consistent with a dysfunctional protein product in the case where a putative negative feedback loop regulates gene expression [21].

**Figure 3.**
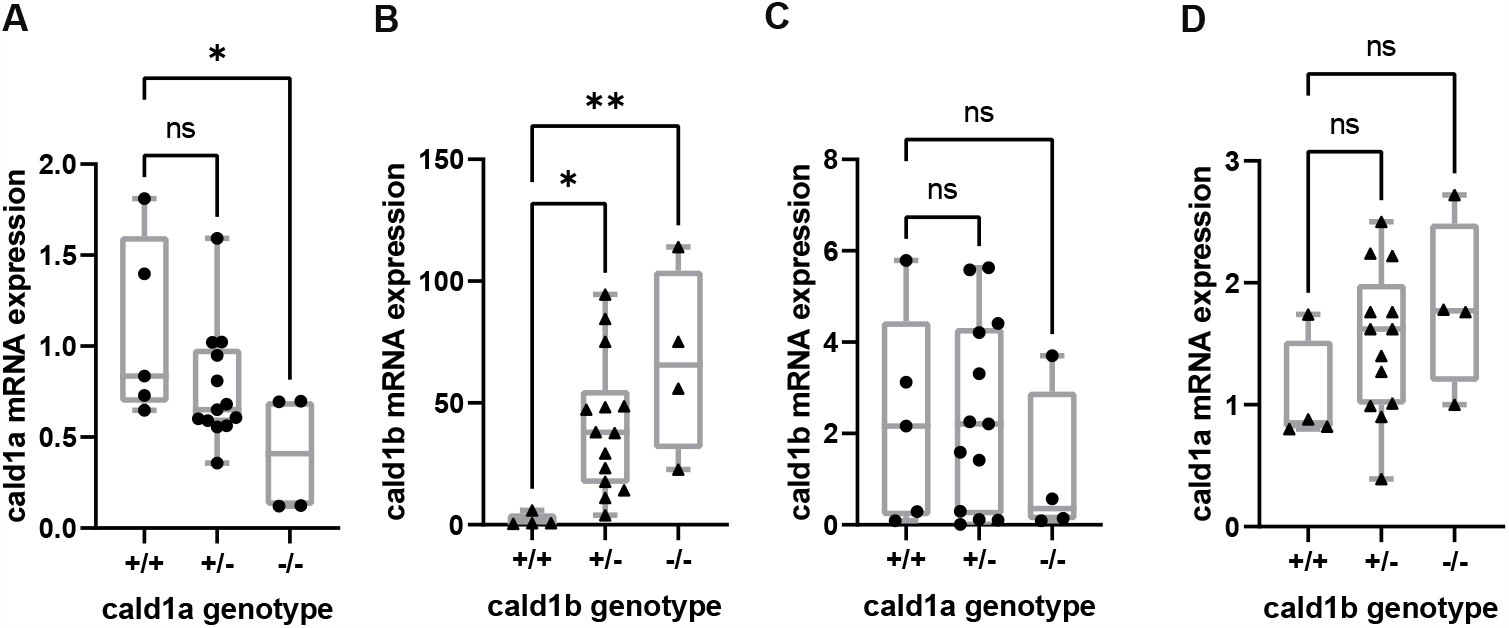
*cald1a* and *cald1b* mutant mRNAs have abnormal expression levels. (A) Relative *cald1a* mRNA expressions in different genotypes of *cald1a* (left) (n_WT_=5, n_(+/−)_=15, and n_(−/−)_=4). (B) Relative *cald1b* mRNA expressions in different genotypes of *cald1b* (right) (n_WT_=4, n_(+/−)_=14, and n_(-/-)_=4) in zebrafish larvae (C) Relative *cald1b* mRNA expressions in different genotypes of *cald1a* (right) (n_WT_=5, n_(+/−)_=13, and n_(−/−)_=4). (D) Relative *cald1a* mRNA expressions in different genotypes of *cald1b* (left) (n_WT_=4, n_(+/−)_=14, and n_(−/−)_=4) in zebrafish larva. ns = not significant, \**p* < 0.05, \*\**p* < 0.01, and *** *p* < 0.001 as determined by ANOVA.

Previous work has indicated that mutant phenotypes could often be alleviated by compensatory mRNA expression from closely related genes upon nonsense-mediated decay of related transcripts [22]. To address this issue, we analyzed *cald1b* mRNA expression in *cald1a* mutants and vice versa. Neither *cald1a* mutants showed compensatory upregulation of *cald1b* mRNA (Fig. 3C) nor the *cald1b* mutants displayed increased compensatory expression of *cald1a* mRNA (Fig. 3D). Despite the lack of compensation response, the *cald1a* and *cald1b* could still have redundant functions and both genes might need to be mutated to see robust phenotypes.

### No clear morphological phenotype for cald1a and cald1b double-mutant embryos

To investigate the effect of simultaneous mutation of both *cald1a* and *cald1b* on the morphology of developing zebrafish embryos, we measured standard length, area of the cephalic region, eye area, and pericardial area using manual segmentation of the brightfield images in ImageJ. No significant difference between the genotypes was detected in the body length, head size, eye size, or pericardium in double-mutant embryos (Fig 4A–E).

**Figure 4.**
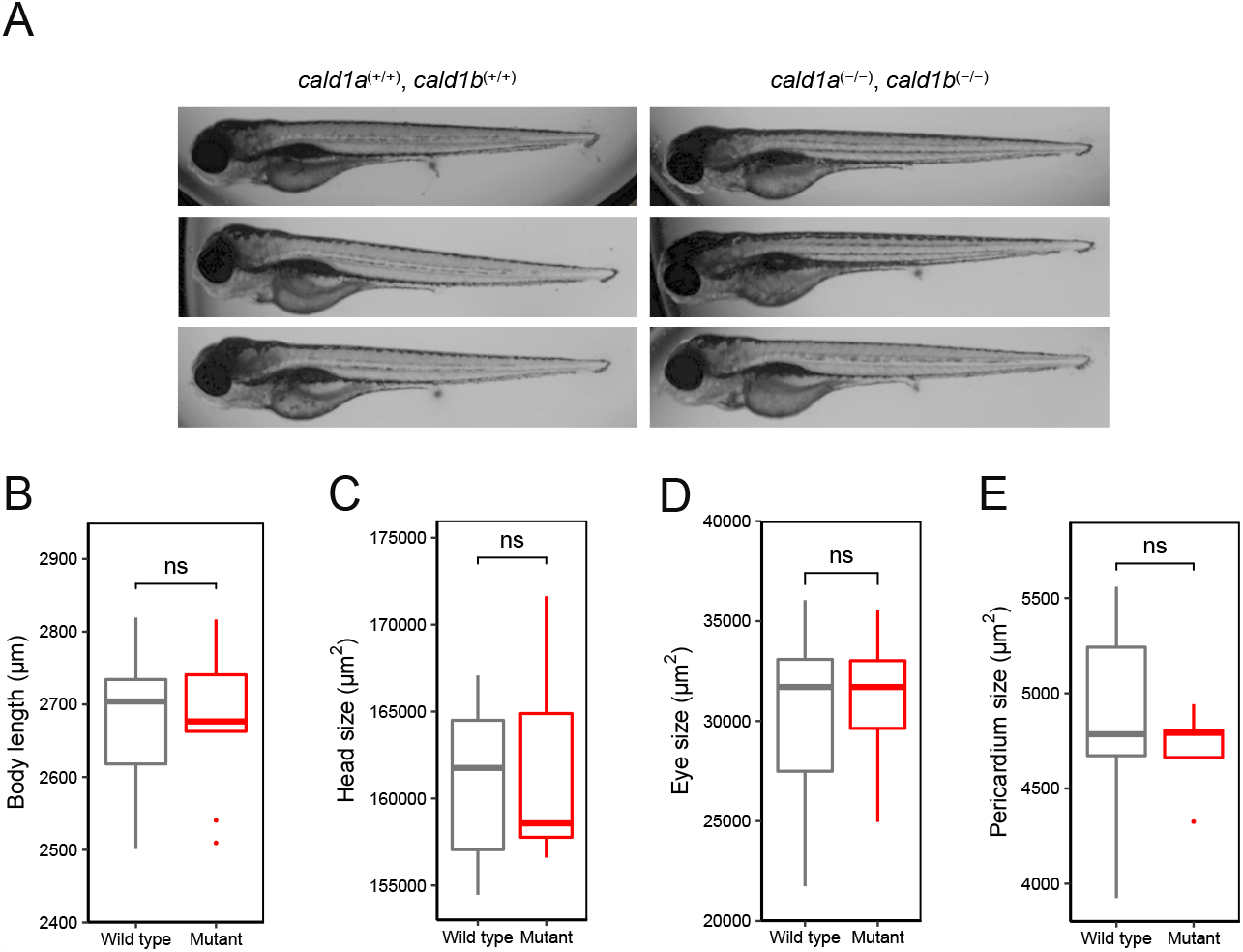
*cald1a, cald1b* double-mutant zebrafish have no significant morphological phenotype. (A) Representative brightfield images of wild type zebrafish larvae *cald1a* (+/+), *cald1b* (+/+) (left) and mutant zebrafish larvae *cald1a* (−/−), *cald1b* (−/−) (right) at 4 dpf. (B–E) Morphological measurements and analyses display no statistically significant changes (ns) in body length, head size, eye size, and in the pericardial area when compared wild type *cald1a* (+/+), *cald1b* (+/+) larvae (n=15) to double-mutant *cald1a* (−/−), *cald1b* (−/−) larvae (n=10).

### Mutation of cald1a and cald1b genes is not lethal for zebrafish larvae

The lack of phenotype could be resulting from loss of strongly affected *cald1*-mutant embryos very early in the development. To identify potential early embryonic lethality, we analyzed the distribution of genotypes in the offspring of double heterozygote parents. The distribution of genotypes followed expected allele frequencies and also the Mendelian inheritance pattern of a dihybrid cross (Fig. 5A, B, C, D and E), indicating that there was no early embryonic lethality caused neither by single *cald1a* or *cald1b* mutation, nor by double homozygous mutation in *cald1a* and *cald1b*.

**Figure 5.**
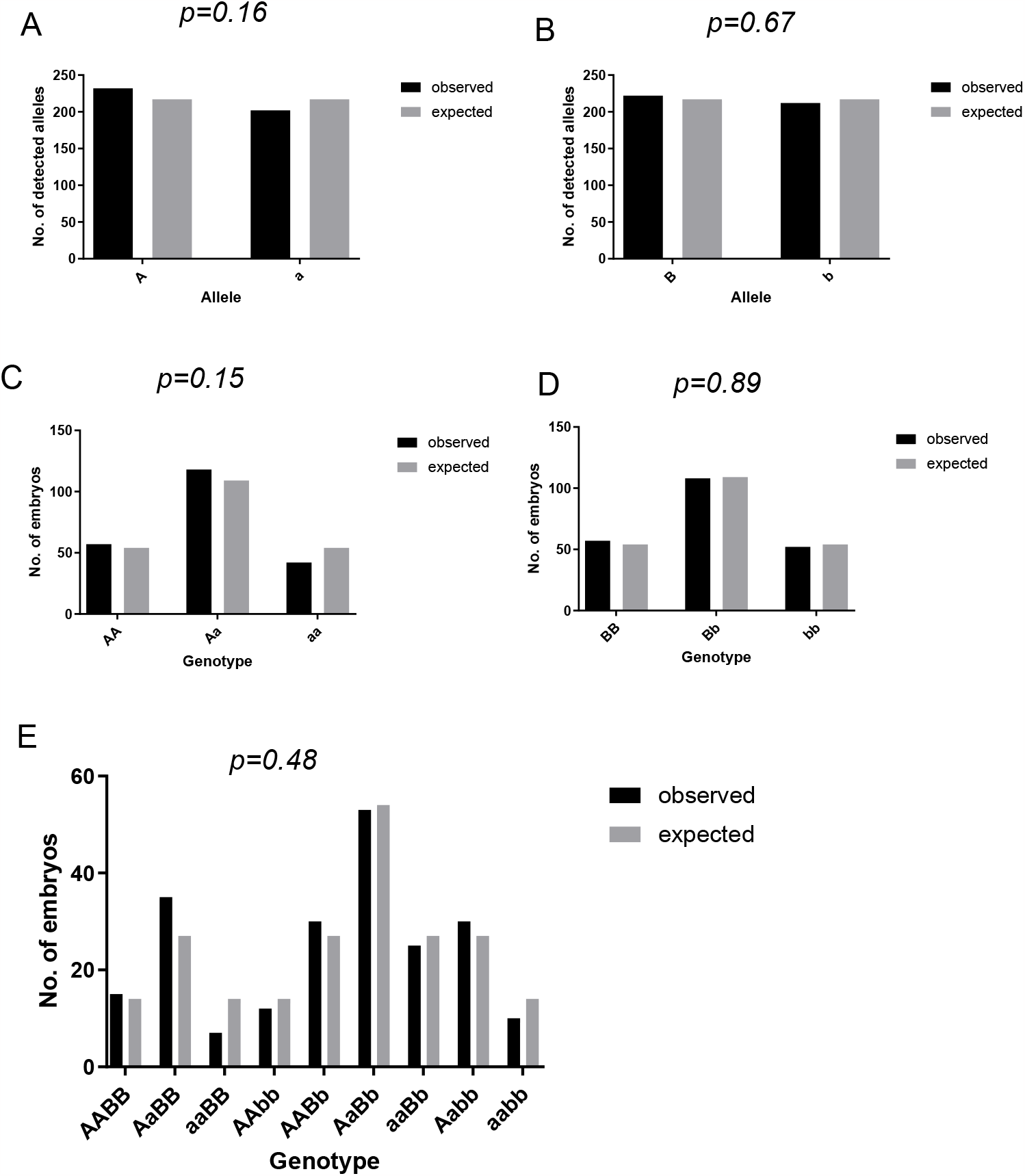
Mutation of cald1a and cald1b genes is not lethal for zebrafish larvae. Analyses of zebrafish allele frequencies and genotype of offspring. Fish heterozygous for both *cald1a* and *cald1b* mutation were incrossed in a dihybrid cross. All embryos were genotyped and observed counts (n = 217) were statistically compared to expected counts using binomial (A and B) or Chi-square (C, D, and E) test. (A) Count of detected alleles of *cald1a* wild-type (marked with A) and sa24667 mutant (a) allele. (B) Count of detected alleles of *cald1b* wild-type (marked with B) and sa16974 mutant (b) allele. (C) Distribution of *cald1a* genotypes. (D) Distribution of *cald1b* genotypes. (E) Distribution of *cald1a, cald1b* dihybrid genotypes.

### cald1-mutation has mild effects on the behavior of zebrafish embryos

Many gene effects are not evident at gross morphological level or survival, but have more subtle impact physiological functions of the organism. In zebrafish, the motility of larvae is widely used assay to measure effects on organismal locomotion and behavior [23]. Therefore, we analyzed the embryos from a *cald1a, cald1b* dihybrid cross for the ability to move and respond to stimulus.

The analysis of fish for the light-response indicated that the *cald1a, cald1b* double mutant fish displayed weaker responses to alternating light-dark illumination (Fig. 6A). This indicated that the *cald1a, cald1b* double mutant fish had mild neurological defect. *cald1a* gene mutation alone caused a milder phenotype (Fig. 6B), but the simultaneous mutation of also *cald1b* gene potentiated this effect. Mutation in *cald1b* gene did not have an effect (Fig 6C). To further analyze if the effect was on visual detection of changes in lighting or the actual ability to move, we stimulated embryos with epileptogenic compound pentylenetetrazol (PTZ). The addition of PTZ induced embryo motility, and prevented light-responses. The *cald1a, cald1b* double mutants, however, did not differ from controls in this setting (Fig. 6D). This implied that the effect of *cald1* mutation was occurring at visual perception and neurological signal processing levels rather than due to better contraction capabilities of muscles.

**Figure 6.**
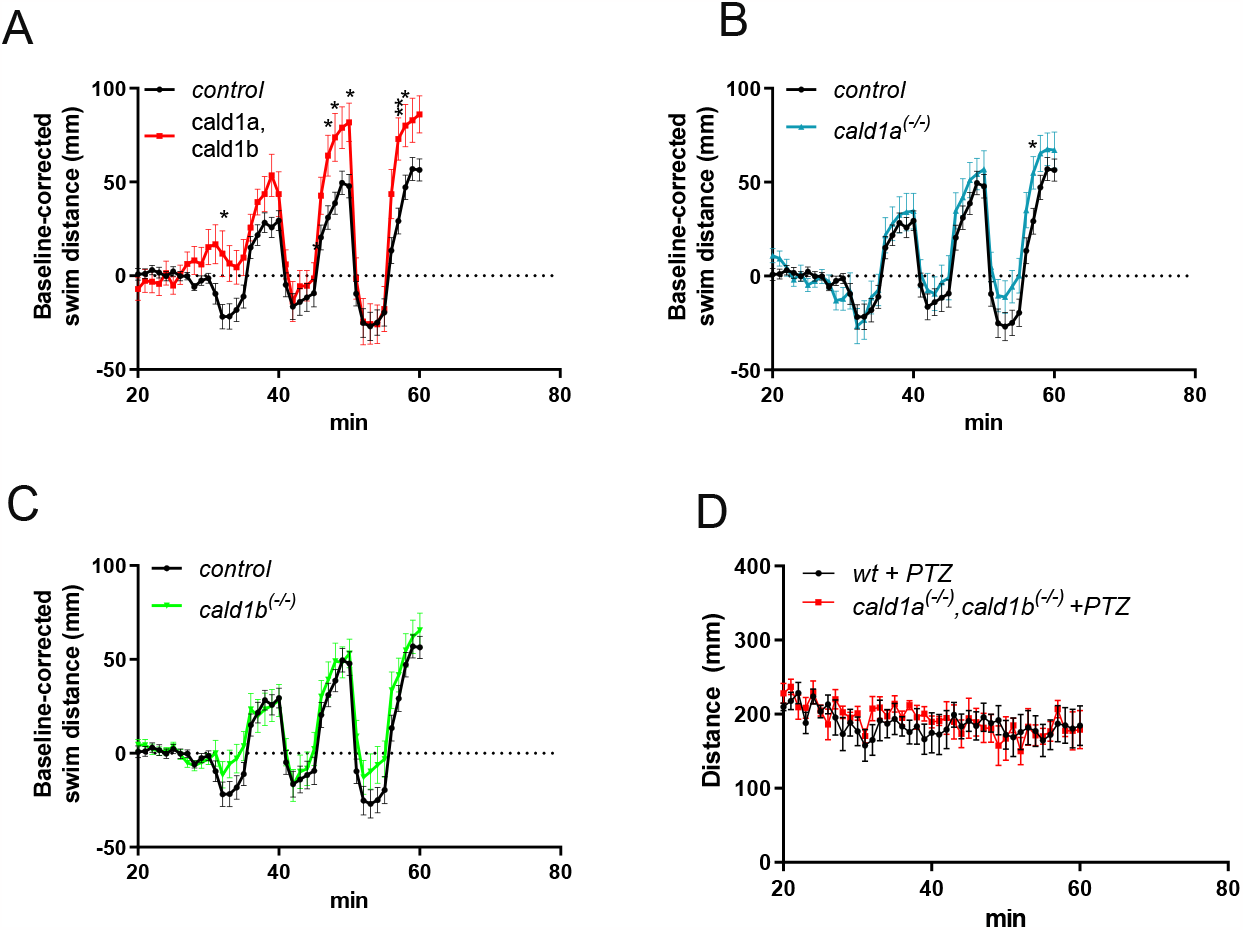
Cald1 mutation has mild locomotory phenotype in zebrafish larvae. Analyses of behavior and locomotion of *cald1a* and *cald1b* mutants in 96-well assay using DanioVision instrument. After the adaptation phase, an alternating light-dark cycles (5min light, 5min darkness) were used to stimulate movements. Some genotypes were pooled to increase statistical power. A) *control* (n= 52, genotypes *wt; cald1a*^*+/-*^ *and cald1b*^*+/-*^) vs *cald1a, cald1b* double-mutants (n=13, genotypes, *cald1a*^-/-^, *cald1b*^-/-^), B) *control* vs *cald1a* mutants (n=24, genotypes cald1a^-/-^ and cald1a^-/-^, cald1b^+/-^), C) *control* vs *cald1b* mutants (n= 38, genotypes cald1b^-/-^ and cald1b^-/-^,cald1a^+/-^), D) Stimulation of wt (n=5) or *cald1a,cald1b* mutants (n=5) with pentylene tetrazolium (PTZ). In A-C the same control data are shown as reference.

## Discussion

By constructing a *cald1a*-*cald1b*-mutant zebrafish model, we demonstrate that *cald1* mutations are not lethal, and no difference in the phenotype between the wild type and the double-mutant is present during early development. Our phenotype analysis was performed on dpf 4; therefore, potential later changes in the phenotype cannot be excluded. Previous work shows *cald1* knockdown in a zebrafish morphant model that presents with serious defects in vasculogenesis, angiogenesis and cardiac organogenesis observed at dpf 1.5–5 [24,25]. More recent zebrafish works put forward considerable criticism for morpholino studies, and therefore confirmation with an appropriate mutant model is recommended [26]. Morphant phenotypes are often more severe and can differ from mutant phenotypes for various reasons, including frequent off-target effects [26]. It is also possible that the mutant lines in our study are hypomorphic and not fully amorphic alleles.

In our studies, we observed effects of *cald1* mutation on light-dark cycle induced motility. As *cald1a* was clearly expressed in retinal and retinal progenitor cells but not in skeletal muscles, and there was no differences upon pharmacological stimulation with PTZ, it seems plausible that the effect on light-dark cycle induced responses occurs at the visual perception and signal processing level. The embryos were still qualitatively responding correctly to the visual stimulus, but the magnitude of the response was affected. This data indicates that the mutations in *cald1a* and *cald1b* have functional effect. This doesn’t fully reject possibility for hypomorphic mutation, but nevertheless indicates that at least some inhibition of cald1 could be tolerated reasonably well.

In the published knockout mice lacking both caldesmon isoforms (l-caldesmon and h-caldesmon), the mice die perinatally with an unresolved umbilical hernia [27]. However, in a model with homozygous loss of h-caldesmon, the loss is shown not to be lethal [28,29]. Another mouse model with mutations targeting a functional domain in both isoforms is lethal as a homozygote but reproduces normally as a heterozygote [30]. The severe cardiovascular defects are not described in *Cald1*-deficient mice, although the underlying causes of pre- and postnatal lethality are not fully understood [28–30]. More comprehensively the postnatal relevance of *Cald1-*deficiency could be evaluated in future studies by using conditional mouse models. Depletion of *cald1* in a xenopus morpholino model leads to severe cartilage defects due to reduced neural crest migration, suggesting its critical function in normal neural crest migration in xenopus [31]. Taken together, *Cald1* may have critical roles in cell migration during early development. However, our results show no strong phenotype in *cald1*-mutated zebrafish larvae, which suggests tolerability towards caldesmon inhibition to a certain degree in healthy tissues, which is a prerequisite for caldesmon being a therapeutic target.

## Acknowledgements

We thank Zebrafish Core, Cell Imaging Core and Finnish Functional Genomics Centre (all in Turku Bioscience Centre and supported by Biocenter Finland) for services, instrumentation, and consultations. We thank Minna Santanen for excellent technical assistance.

## Funding

This work was supported by grants from Academy of Finland, Finnish Medical Foundation, Finnish Cancer Foundations, Turku University Foundation, Turku University Hospital, TYKS Foundation, Finnish Cultural Foundation, and Turku Doctoral Programme of Molecular Medicine (TuDMM).

## Contributions

VV, KP, IP, and MS designed the experiments; KP and IP carried out the zebrafish experiments; VV, KP, SN, and IP extracted measurements from the zebrafish; VV, KP, and IP prepared the figures; IP, KP, VV and MS prepared the original draft; VV, KP, IP, and MS reviewed and edited the article; IP and MS supervised the study; MS acquired funding; All authors have read and agreed to the published version of the manuscript.

## Conflict of Interests

Authors declare no competing financial interests.

## Notes

### Competing Interest Statement

The authors have declared no competing interest.

